# Hippocampal and large-scale functional connectivity reorganization following freediving training, and relationships with episodic memory

**DOI:** 10.64898/2026.09.24.753809

**Authors:** Julia Micaux, Abir Troudi Habibi, Franck Mauconduit, Fawzi Boumezbeur, Catherine Chiron, Marion Noulhiane

## Abstract

The hippocampus is particularly vulnerable to hypoxia, yet freedivers, who repeatedly undergo voluntary breath-hold hypoxia, do not appear to show the memory impairments commonly reported after involuntary hypoxic exposure. We investigated whether freediving training is associated with functional connectivity reorganization at the whole-brain and hippocampal levels, and whether hippocampal connectivity relates to preserved episodic memory performance. Seventeen male recreational freedivers underwent fMRI before and after a 7-month standardized training period, and 20 matched aerobic-trained controls were assessed at baseline. Functional connectivity analyses examined (i) whole-brain ROI-to-ROI connectivity, (ii) seed-to-voxel connectivity from the bilateral hippocampus and selected cortical seeds, and (iii) associations between hippocampal connectivity and episodic memory. Compared with controls, freedivers showed distinct connectivity patterns during apnea and rest. After training, whole-brain analyses revealed reorganization involving salience, frontoparietal, sensorimotor, visual, and cerebellar systems. Hippocampal analyses showed reduced connectivity with sensorimotor and default mode regions, alongside increased connectivity with cerebellar and visual areas, with stronger effects after training. In freedivers, specific post-training hippocampal connectivity was associated with episodic memory performance. These findings suggest that repeated voluntary hypoxia during freediving training is associated with selective functional reorganization of hippocampal and large-scale brain networks. This pattern may reflect adaptive neuroplasticity linked to preserved episodic memory under intermittent hypoxic exposure.

**Key points:**

- Freedivers exhibit selective hippocampal functional connectivity differences during voluntary intermittent breath-hold hypoxia, including reduced coupling with sensorimotor and default mode regions and increased coupling with cerebellar areas.
- A 7-month training period is associated with large-scale network reorganization involving salience, frontoparietal, sensorimotor, visual, and cerebellar systems.
- Post-training hippocampal connectivity is associated with preserved episodic memory performance, consistent with adaptive functional reorganization under voluntary intermittent hypoxia.

## Introduction

The hippocampus plays a central role in episodic memory, supporting the encoding, storage, and retrieval of experiences within their spatial and temporal contexts. A key process that supports this function is pattern separation, which allows the discrimination of similar distinct events and depends primarily on hippocampal circuitry, particularly the dentate gyrus and CA3 subfields (Castillon et al., 2018). Since the HM case (Scoville and Milner, 1957), hippocampal damage has been consistently associated with profound episodic memory disorders.

Among factors affecting hippocampal integrity, hypoxia represents a major challenge. Involuntary hypoxia, arising from pathological or environmental conditions, is known to disrupt brain function, with the hippocampus being particularly vulnerable due to its high metabolic demand and complex vascularization (Spallazzi et al., 2019), making it one of the first structures affected by oxygen deprivation in multiple clinical contexts (Lu et al., 2025; McKeown et al., 2025). Clinical and experimental studies have reported hippocampal atrophy, impaired neurogenesis, and reduced synaptic plasticity following hypoxic exposure; often accompanied by deficits in memory, attention and executive functions (Khuu et al., 2019; Li and Fei, 2013; Lu et al., 2025; Olaithe et al., 2018; Pfister et al., 2023; Zhang et al., 2022).

Resting-state functional connectivity (rs-FC) offers a non-invasive tool to study how large-scale brain networks are altered under hypoxic conditions. Previous rs-fMRI studies of involuntary hypoxia have reported widespread reductions in FC and network disorganization, particularly involving hippocampal networks and the default mode network (DMN) (Bakker et al., 2023; Micaux et al., 2025b; Song et al., 2018). These alterations are typically associated with impaired cognitive performance, such as learning and memory, attention, processing speed, and executive control (McKeown et al., 2025; Wang et al., 2022).

In contrast, freedivers (FD) represent a unique model of repeated voluntary intermittent hypoxia in healthy individuals. Despite frequent exposure to low oxygen levels during breath-hold training, freedivers do not appear to exhibit the cognitive impairments observed in clinical hypoxia (Allinger et al., 2024; Ridgway and McFarland, 2006). Our previous work has shown preserved hippocampal structure and episodic memory performance in this population (Micaux et al., 2025a), suggesting that adaptive mechanisms may support neural resilience. Freediving also combines hypoxic exposure with exercise training, both of which are known to modulate brain plasticity. Exercise has consistently been associated with functional reorganization of large-scale networks and enhanced hippocampal function (Heller-Wight et al., 2023; Li et al., 2023; Raichlen et al., 2016; Suwabe et al., 2018).

In FD, a study by Annen et al. (2021) suggests a similar pattern of adaptation. Increased FC was observed during apnea in subcortical regions (bilateral thalamus, caudate, right amygdala) and in cortical territories involving the DMN and executive control networks. In parallel, decreased FC was mainly found in sensorimotor regions but also in temporal, parietal, insular, and occipital cortices. These findings suggest that freediving promotes functional optimization, enhancing control and introspection while reducing sensory interference, similar to patterns observed in other adaptive contexts such as meditation or exercise training. Additionally, intermittent hypoxia protocols have been proposed to induce neuroprotective effects under specific conditions (Damgaard et al., 2023). However, the functional mechanisms underlying these adaptations in freedivers remain largely unexplored, particularly at the level of hippocampal connectivity.

The present study addresses this gap by examining whole brain and hippocampal functional connectivity in male freedivers before and after a 7-month training period. We expected that: (1) Freedivers would show FC patterns consistent with large-scale network reorganization; (2) Hippocampal FC would exhibit selective changes involving key cognitive and sensorimotor networks; (3) Hippocampal FC would be associated with episodic memory performance, particularly after training.

## Material and Methods

### 1. Participants

Data acquisition was conducted in accordance with the regulations of an accredited Ethical Committee board (CPP 2022-A01030-43). Written informed consent was obtained from all participants before inclusion. Thirty-seven men participated in this study, including 17 FD matched to 20 NC aged from 27-55 years, and recruited between 2022 and 2024 from swimming pools (Ile de France, FRANCE).

The FD group included experienced recreational FD, all affiliated with a freediving club to ensure consistent and structured training. They underwent a 7-month standardized freediving training (Lemaître, 2015; Micaux et al., 2025a), with data collected before (FD_t0_) and after (FD_t1_) training. The NC group consisted of age- and sex-matched participants with comparable socio-cultural backgrounds (Table 1). Although not engaged in freediving, they maintained a similar weekly exercise training load (∼5 hours per week) through aerobic sports. NC participants underwent a single MRI at t_0_ and completed the memory task at both t_0_ and t_1_.

**Table 1.** Demographic data (FD and NC)

| Characteristics | Freedivers (FD) | Controls (NC) |
| --- | --- | --- |
| n | 17 | 20 |
| sex | H | H |
| age* | 47.9 ± 4.8 | 44.8 ± 9 |
| years of study* | 16.9 ± 1.7 | 16.5 ± 2.4 |
| Training load | ~ 5 h/week<br>(aerobic & anaerobic) | ~ 5h/week<br>(aerobic) |

\* Data are means ± standard deviation.

### 2. Episodic memory assessment

Episodic memory was assessed using the HippoPS task (Micaux et al., 2025a), an ecological and validated tool specifically designed to study PS. The task includes two phases: encoding and recognition (after a 15-minute delay). During encoding, participants watch videos showing 28 pictures of a location associated with a gesture. During recognition, they categorize each photo as: i. ‘*identical*’ (same gesture and location), ii. ‘*similar*’ (same gesture or same location), or iii. ‘*new*’ (different gesture and location). These scores were correlated with hippocampal FC (Figure 1). Global intelligence was also assessed, revealing no cognitive deficits in either group (NC or FD), with comparable scores between cohorts (Micaux et al., 2025a).

**Figure 1:**
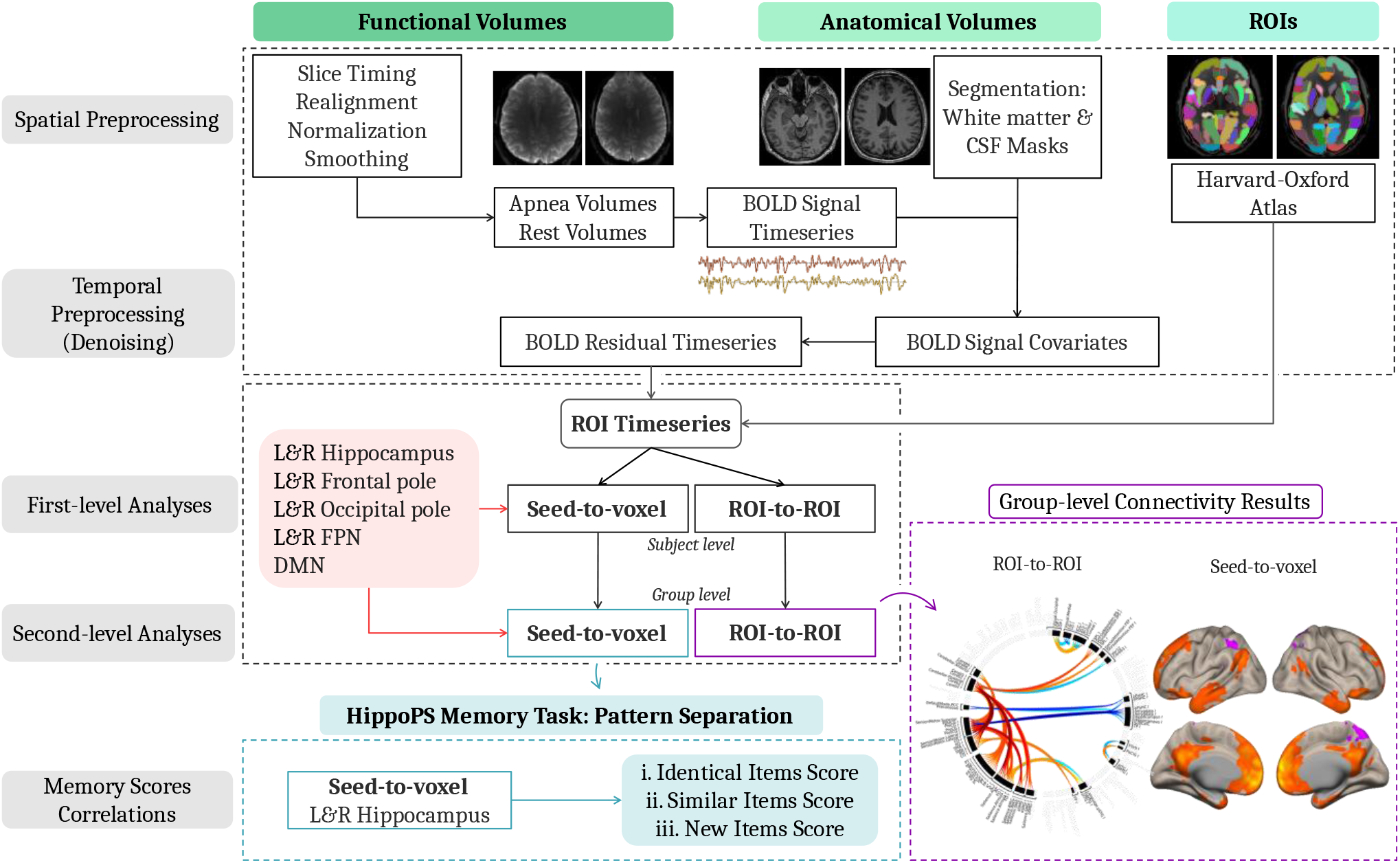
Schematic representation of acquisitions, preprocessing pipeline, and analysis using the CONN toolbox on MATLAB. L&R: Left and right.

### 3. Rs-fMRI acquisitions

Cerebral MRI was performed at 3T (Magnetom Prisma, Siemens Healthineers), including acquisition of T_1_ and T_2_-weighted images of the hippocampus (Figure 1) (Table 2).

**Table 1.** MRI acquisition parameters.

| MRI modality | Sequence | TR (ms) | TE (ms) | Voxel size (mm) | Slices | Duration |
| --- | --- | --- | --- | --- | --- | --- |
| Anatomical MRI | T<sub>1</sub>-weighted (3D MP2RAGE) | 5000 | 2.98 | 1.0 x 1.0 x 1.0 | 176 | 8' 22" |
|  | T<sub>2</sub>-weighted (Hippocampal slices) | 9600 | 83.0 | 0.5 x 0.5 x 1.0 | 60 | 6' 46" |
| Functional MRI | Spin-Echo Fieldmap x2 (AP-PA) | 3510 | 55.4 | 1.7 x 1.7 x 1.8 | 68 | 0' 29" |
|  | BOLD fMRI (EPI multiband) | 2500 | 26.0 | 1.7 x 1.7 x 1.8 | 68 | 3' 30" / run |

Each freediver completed two resting-state fMRI (rs-fMRI) sessions before (FD_t0_) and after (FD_t1_) training. We acquired 4 resting-state fMRI runs, each lasting 3 minutes and 30 seconds. During each run, participants underwent a 2-minute apnea period followed by 1 minute and 30 seconds of regular breathing. A 1-minute rest period was included between runs. Throughout each run, participants viewed an abstract movie adapted from (Vanderwal et al., 2015), a low-demand paradigm developed to improve compliance while preserving the intrinsic organization of large-scale resting-state networks. Apnea instructions were superimposed onto this paradigm to standardize breath-hold timing across participants. The abstract movie began with a countdown displaying the message “Apnea in 10s” down to “Apnea” at 0. After 2 minutes, the message “End of apnea” was displayed.

### 4. Rs-fMRI preprocessing

Rs-fMRI data were preprocessed using the default preprocessing pipeline of the CONN Toolbox (version 20b) (Figure 1). Functional volumes were first realigned and unwrapped to correct for subject motion and geometric distortions. Outlier scans were identified using the ART toolbox, based on both framewise displacement thresholds (0.9 mm) and global signal Z-value thresholding (95th percentile), with additional detection of volumes falling outside the MNI space. Functional and anatomical images were segmented and normalized to MNI space. A spatial smoothing step was applied using an 8 mm FWHM Gaussian kernel to reduce anatomical variability and enhance signal-to-noise ratio. During denoising, temporal pass-band filtering (0.008–0.09 Hz) was performed to retain low-frequency resting-state fluctuations, and physiological noise was removed using aCompCor by regressing out confounding signals from white matter, CSF, and motion parameters. Visual inspection and quality control were conducted at each stage of preprocessing.

### 5. Statistical analysis

All statistical analyses were performed using CONN Toolbox implemented in MATLAB (MathWorks, USA), with a significance threshold set at p < 0.05, false discovery rate (FDR) corrected at the cluster level (Figure 1).

For the FD group, each of the four rs-fMRI runs was temporally divided into two separate runs, apnea and rest, based on each participant’s individual breath-hold duration. This ensured physiological accuracy, as some participants voluntarily ended apnea before the 2-minute instruction. We analyzed FC of the bilateral hippocampus and four additional regions of interest (ROIs) selected a priori based on vulnerability to hypoxia and their involvement in cognitive control networks (Micaux et al., 2025b): the frontal pole, the frontoparietal network, the default mode network (DMN), and the occipital pole.

#### First level analysis: Connectivity analyses

We first performed whole-brain ROI-to-ROI FC analyses to assess global network-level reorganization. Next, seed-to-voxel analyses were conducted using each ROI as a seed to evaluate the connectivity with every voxel across the brain. All analyses were conducted using the Harvard-Oxford atlas, which provides whole-brain coverage. Connectivity matrices and seed-based connectivity maps were generated separately for each hemisphere.

#### Second level analysis: Comparisons

Group- and condition-level comparisons were conducted using CONN’s second-level general linear model framework. The following contrasts were computed to examine the effects of apnea, training and group differences: (1) FDt_0apnea_ > FDt_0rest_, (2) FDt_1apnea_ > FDt_1rest_, (3) FDt_0apnea_ > FDt_1apnea_, (4) FDt_0rest_ > FDt_1rest_, and (5) all of the above conditions were also compared with the NC group (FDt_0apnea_ > NC, FDt_1apnea_ > NC, FDt_0rest_ > NC, and FDt_1rest_ > NC). These contrasts yielded condition-specific difference maps and FC matrices, enabling the identification of adaptive training-related changes.

#### Rs-fMRI / memory score correlation analysis

Using the CONN correlation toolbox, we performed correlations between FC of the left and right hippocampus and episodic memory scores obtained from our previous study (Micaux et al., 2025a). Correlations were conducted separately for FD and NC, during apnea and rest, before and after training. A second-level general linear model (GLM) was applied, with memory scores entered as covariates.

#### Multiple comparison correction

All statistical tests were corrected for multiple comparisons using the FDR method at *p* < 0.05.

## Results

### 1. Whole-brain connectivity

Whole-brain ROI-to-ROI analysis revealed significant FC differences between NC and FD during both apnea and rest.

During apnea, FD showed a consistent pattern of cerebellar and salience network hyperconnectivity alongside hippocampal hypoconnectivity. Before training (t_0_), cerebellar hyperconnectivity extended to sensorimotor, dorsal attention, and salience networks, while the hippocampus showed reduced connectivity with the DMN and sensorimotor network. After training (t_1_), this pattern partially reorganized: salience-sensorimotor/dorsal attention and hippocampal-cerebellar hyperconnectivity emerged alongside new sensorimotor-subcortical connections (thalamus, caudate, putamen), though hippocampal-sensorimotor hypoconnectivity persisted (Figure 2).

**Figure 2:**
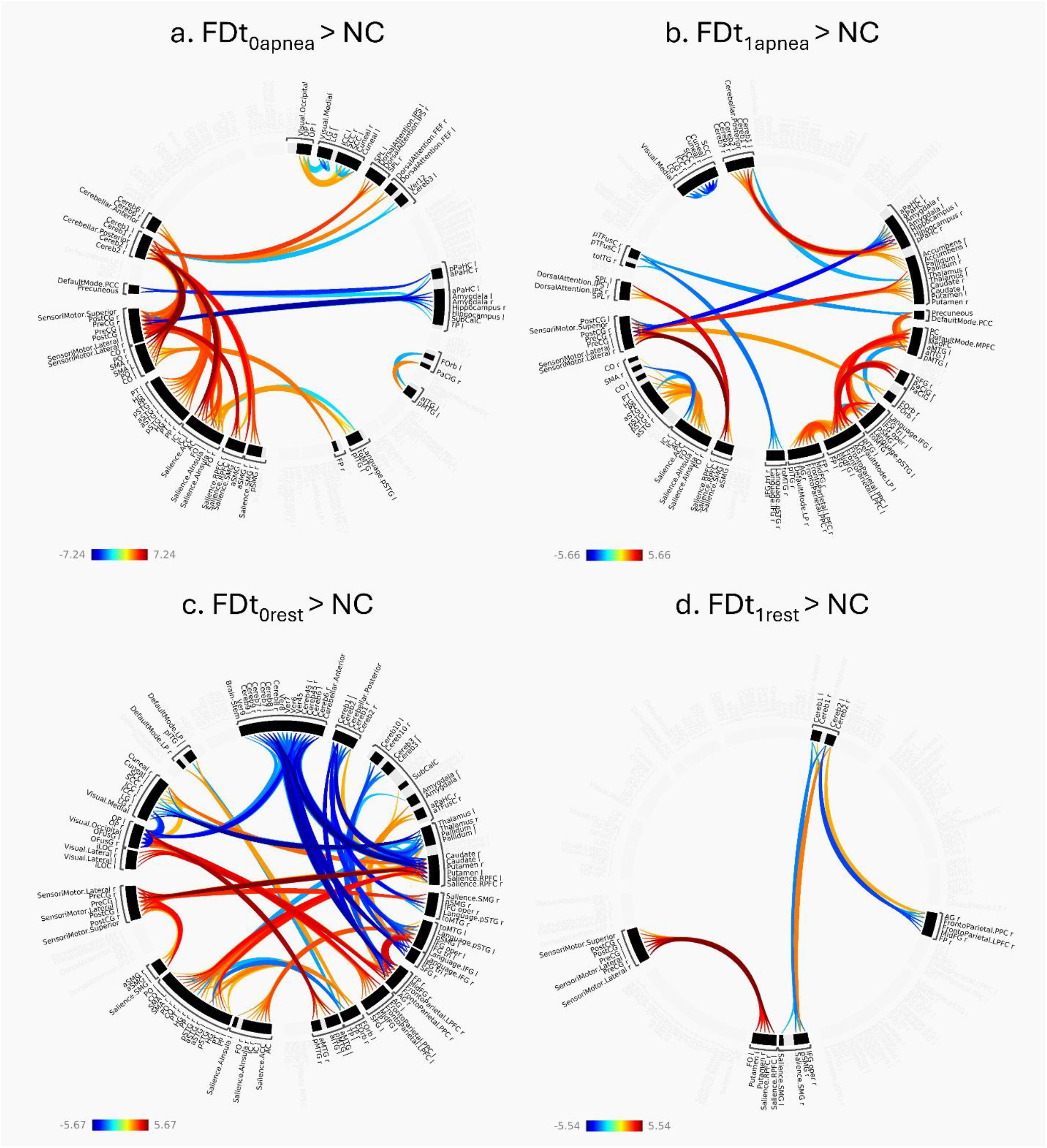
FC matrices showing differences in whole brain hypoconnectivity and hyperconnectivity comparing different conditions (NC versus FD during apnea and rest, before and after training): ROI-to-ROI analysis using CONN toolbox on MATLAB. a. FDt_0apnea_ > NC b. FDt_1apnea_ > NC c. FDt_0rest_ > NC d. FDt_1rest_ > NC. N= 17 FD and 20 NC. FDR correction applied. FD: Freedivers, NC: Controls.

During rest, FD showed a contrasting profile of subcortical and visual hyperconnectivity against broad cerebellar disconnection. Before training (t_0_), the caudate was hyperconnected with sensorimotor, SMA, and STG regions, and the visual network with MTG and FPN, while the cerebellum was prominently hypoconnected across FPN, visual, salience, and language networks. After training (t_1_), cerebellar connectivity partially recovered toward salience and angular gyrus regions, and sensorimotor-putamen/salience hyperconnectivity emerged, yet cerebellar hypoconnectivity with frontoparietal and salience networks persisted (Figure 2).

Comparing apnea to rest before training, FD showed relatively greater cerebellar-salience/sensorimotor hyperconnectivity during apnea, alongside visual-FPN/MTG hypoconnectivity (Figure S1).

### 2. Connectivity in regions of interest

Only significant results are presented, and full statistics are available in Table S1 and S2.

#### 2.1 Hippocampus

Both hippocampi showed reduced connectivity with sensorimotor regions and increased cerebellar connectivity, more pronounced on the left. Before training, the left hippocampus displayed additional hypoconnectivity with default mode regions at rest. Training reinforced cerebellar hyperconnectivity bilaterally while extending sensorimotor hypoconnectivity, particularly at rest. The right hippocampus followed a similar but attenuated pattern, with no significant differences before training at rest (Figure 3).

**Figure 3:**
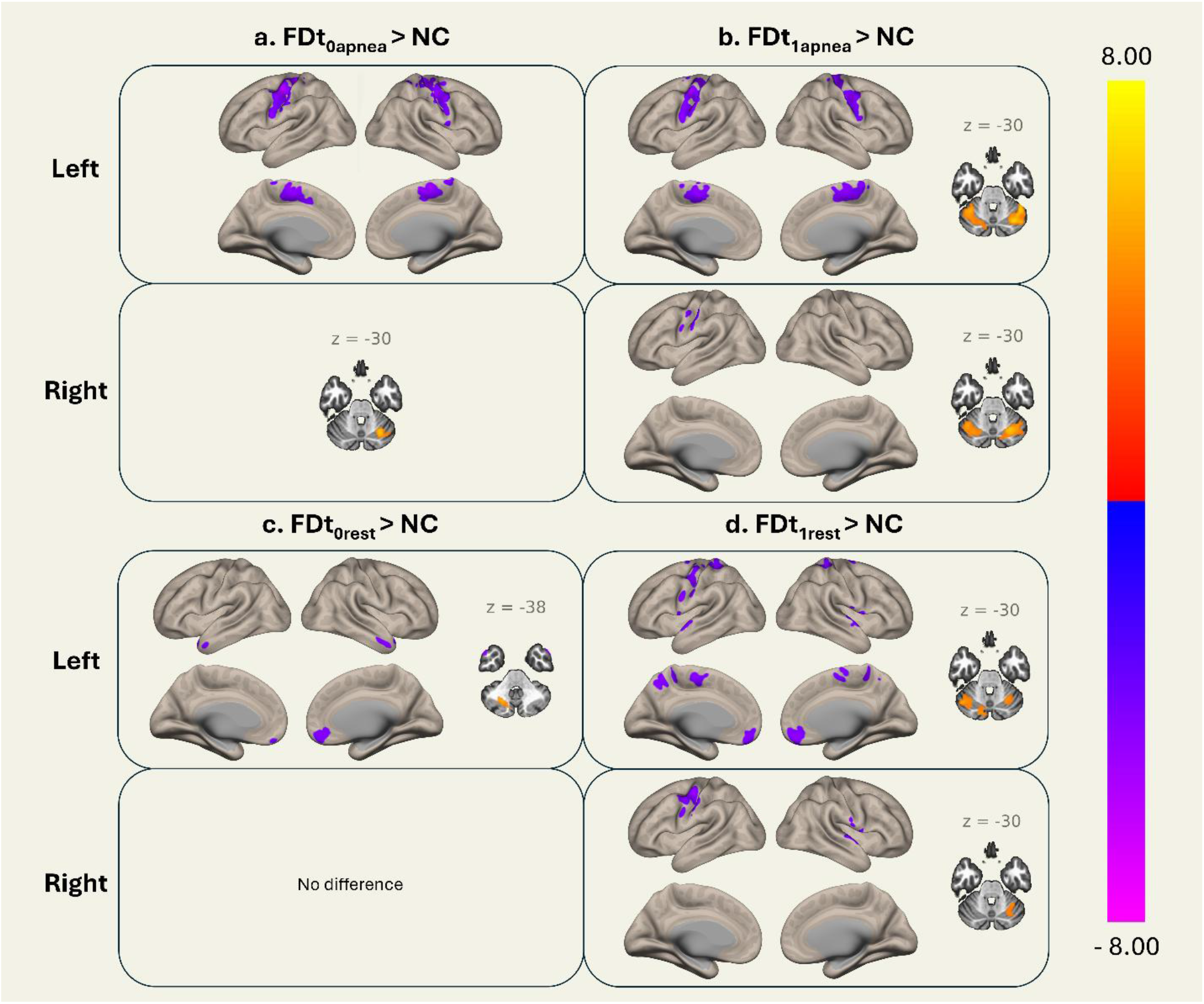
Left and right hippocampal FC differences between conditions (NC versus FD during apnea and rest, before and after training): Seed-to-Voxel analysis using CONN toolbox on MATLAB, FDR correction applied. 2D slices and 3D view. a. FDt_0apnea_ > NC b. FDt_1apnea_ > NC c. FDt_0rest_ > NC d. FDt_1rest_ > NC. n = 17 FD and 20 NC. FD: Freedivers, NC: Controls.

#### 2.2 Default mode network

The DMN showed a focused pattern: training was associated with hypoconnectivity with occipital and cerebellar regions during apnea, while no significant differences were found at rest, before or after training (Figure 4).

**Figure 4:**
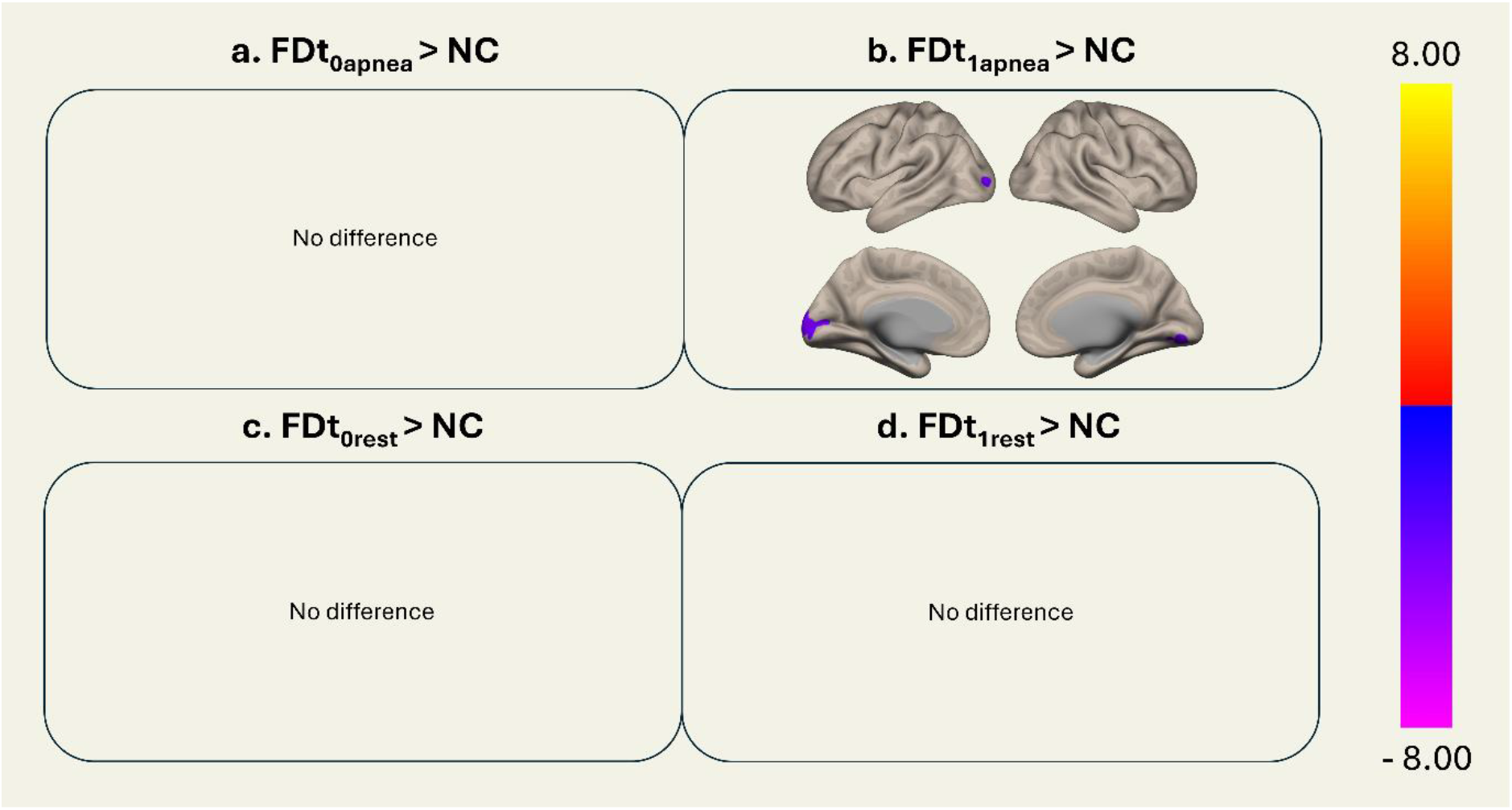
DMN FC differences between conditions (NC versus FD during apnea and rest, before and after training): Seed-to-Voxel analysis using CONN toolbox on MATLAB, FDR correction applied. 2D slices and 3D view. a. FDt_0apnea_ > NC b. FDt_1apnea_ NC c. FDt_0rest_ > NC d. FDt_1rest_ > NC. N= 17 FD and 20 NC. FD: Freedivers, NC: Controls.

#### 2.3 Frontal cortex

Both frontal cortices showed persistent hyperconnectivity with sensorimotor regions (precentral/postcentral gyri, central opercular cortex) during apnea, before and after training, more widespread on the right. At rest, a pattern of cerebellar hypoconnectivity before training shifted after training toward frontal and occipital hypoconnectivity alongside emerging prefrontal hyperconnectivity (Figure 5).

**Figure 5:**
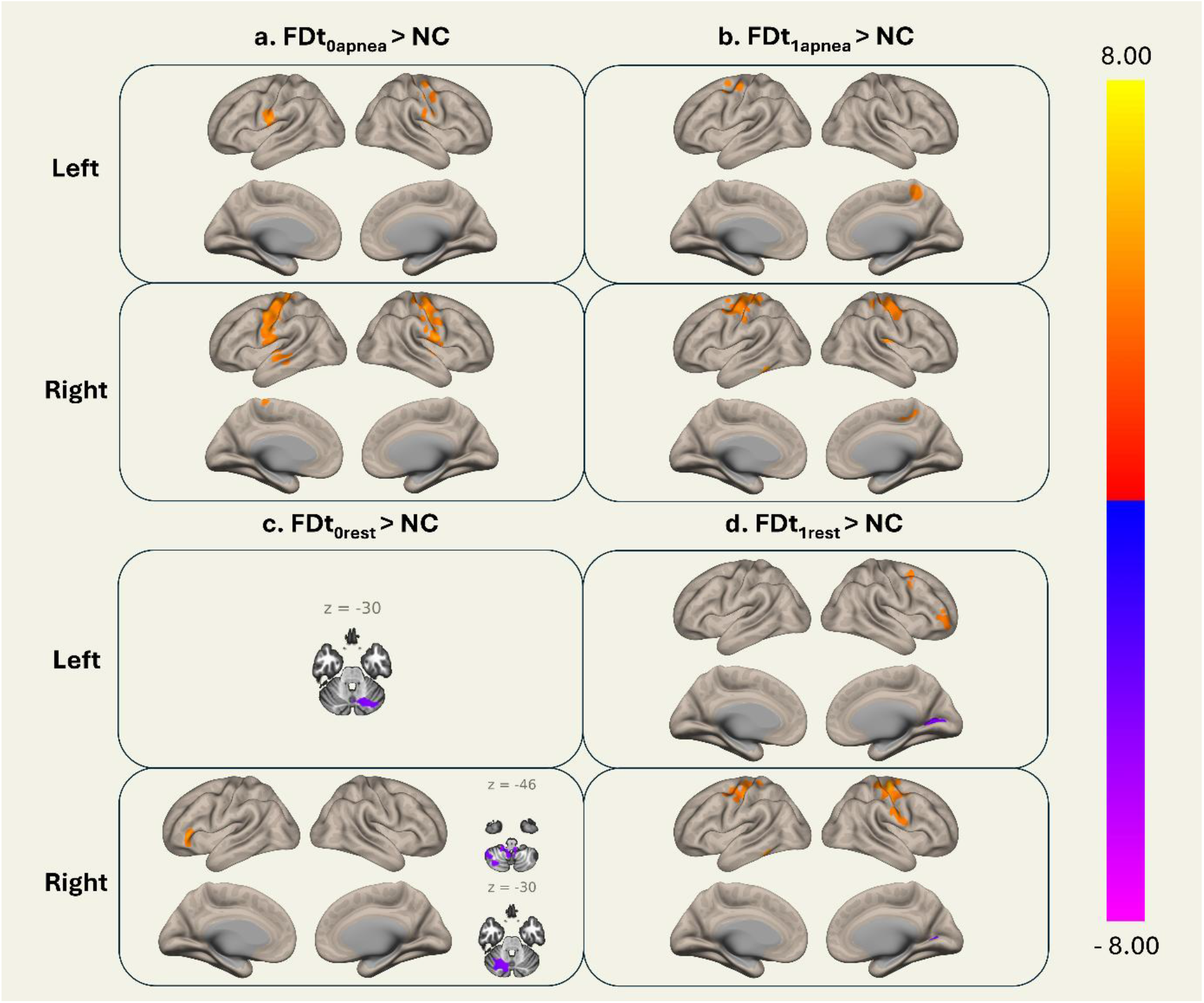
Left and right frontal FC differences between conditions (NC versus FD during apnea and rest, before and after training): Seed-to-Voxel analysis using CONN toolbox on MATLAB, FDR correction applied. 2D slices and 3D view. a. FDt_0apnea_ NC b. FDt_1apnea_ > NC c. FDt_0rest_ > NC d. FDt_1rest_ > NC. N= 17 FD and 20 NC. FD: Freedivers, NC: Controls.

#### 2.4 Frontoparietal network

Both frontoparietal networks showed prominent hyperconnectivity with sensorimotor regions (precentral/postcentral gyri, SMA) during apnea and rest, more extensive on the right. Training amplified this pattern, extending hyperconnectivity to cingulate and insular regions. Cerebellar hypoconnectivity observed before training at rest was largely replaced after training by thalamic and occipital hypoconnectivity (Figure 6).

**Figure 6:**
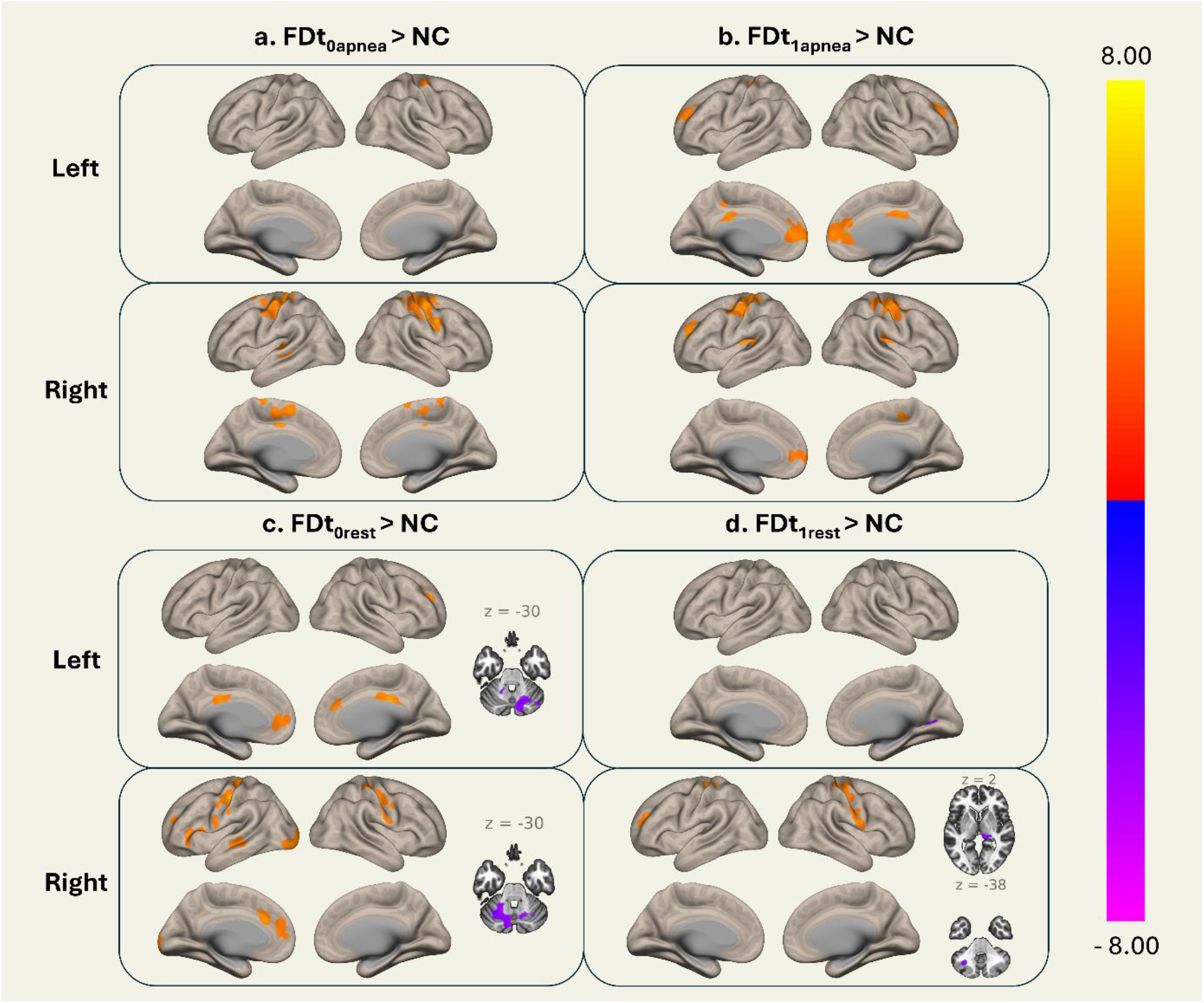
Left and right frontoparietal FC differences between conditions (NC versus FD during apnea and rest, before and after training): Seed-to-Voxel analysis using CONN toolbox on MATLAB, FDR correction applied. 2D slices and 3D view. a. FDt_0apnea_ > NC b. FDt_1apnea_ > NC c. FDt_0rest_ > NC d. FDt_1rest_ > NC. N= 17 FD and 20 NC. FD: Freedivers, NC: Controls.

#### 2.5 Occipital cortex

Both occipital cortices showed a before/after training asymmetry: before training, widespread hypoconnectivity with visual and cerebellar regions coexisted with extensive hyperconnectivity with temporal, parietal, and frontal regions, predominantly at rest. After training, these differences largely disappeared, suggesting a normalization of occipital connectivity. During apnea, effects were more limited and confined to frontal and cingulate hypoconnectivity (Figure 7).

**Figure 7:**
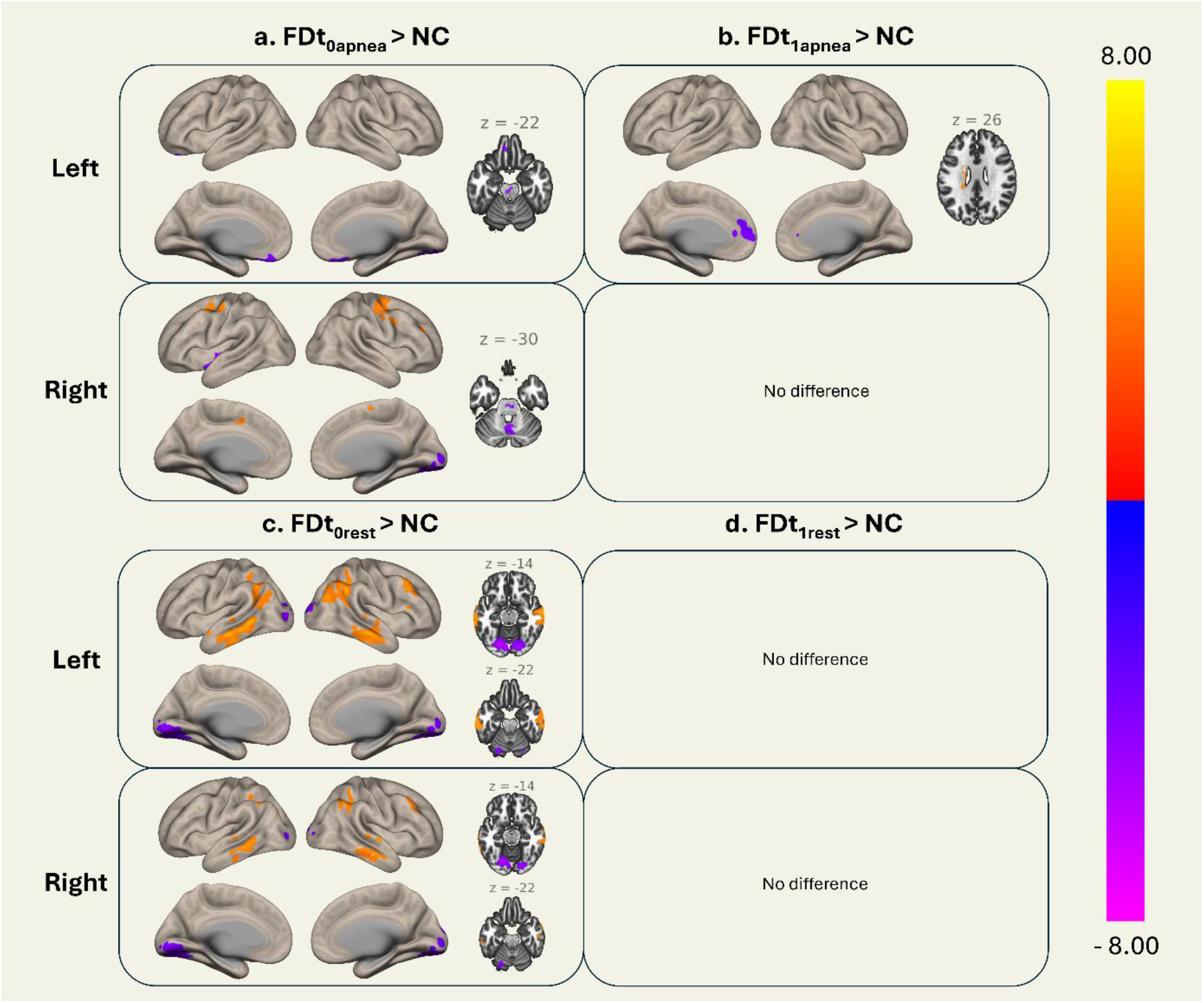
Left and right occipital FC differences between conditions (NC versus FD during apnea and rest, before and after training): Seed-to-Voxel analysis using CONN toolbox on MATLAB, FDR correction applied. 2D slices and 3D view. a. FDt_0apnea_ NC b. FDt_1apnea_ > NC c. FDt_0rest_ > NC d. FDt_1rest_ > NC. N= 17 FD and 20 NC. FD: Freedivers, NC: Controls.

### 3. Episodic memory correlations

#### 3.1 Left hippocampus

##### FD group, rest after training

We found a significant positive correlation between EM scores of ii. *similar* items and the FC of the left hippocampus with the right cerebellum (*r* = 0.89, p<0.001, n = 17) and the left hippocampus (*r* = 0.88, p = 0.044, n = 17). We also found a significant positive correlation between EM scores of iii. *new* items and the FC of the left hippocampus with the right cerebellum (*r* = 0.90, p<0.001, n = 17). No other significant correlations were found in the other conditions.

##### NC group

We found a significant positive correlation between EM scores of ii. *similar* items and the FC of the left hippocampus with the frontal medial cortex and the paracingulate gyrus (*r* = 0.79, p = 0.045, n = 20) (Figure 8).

**Figure 8:**
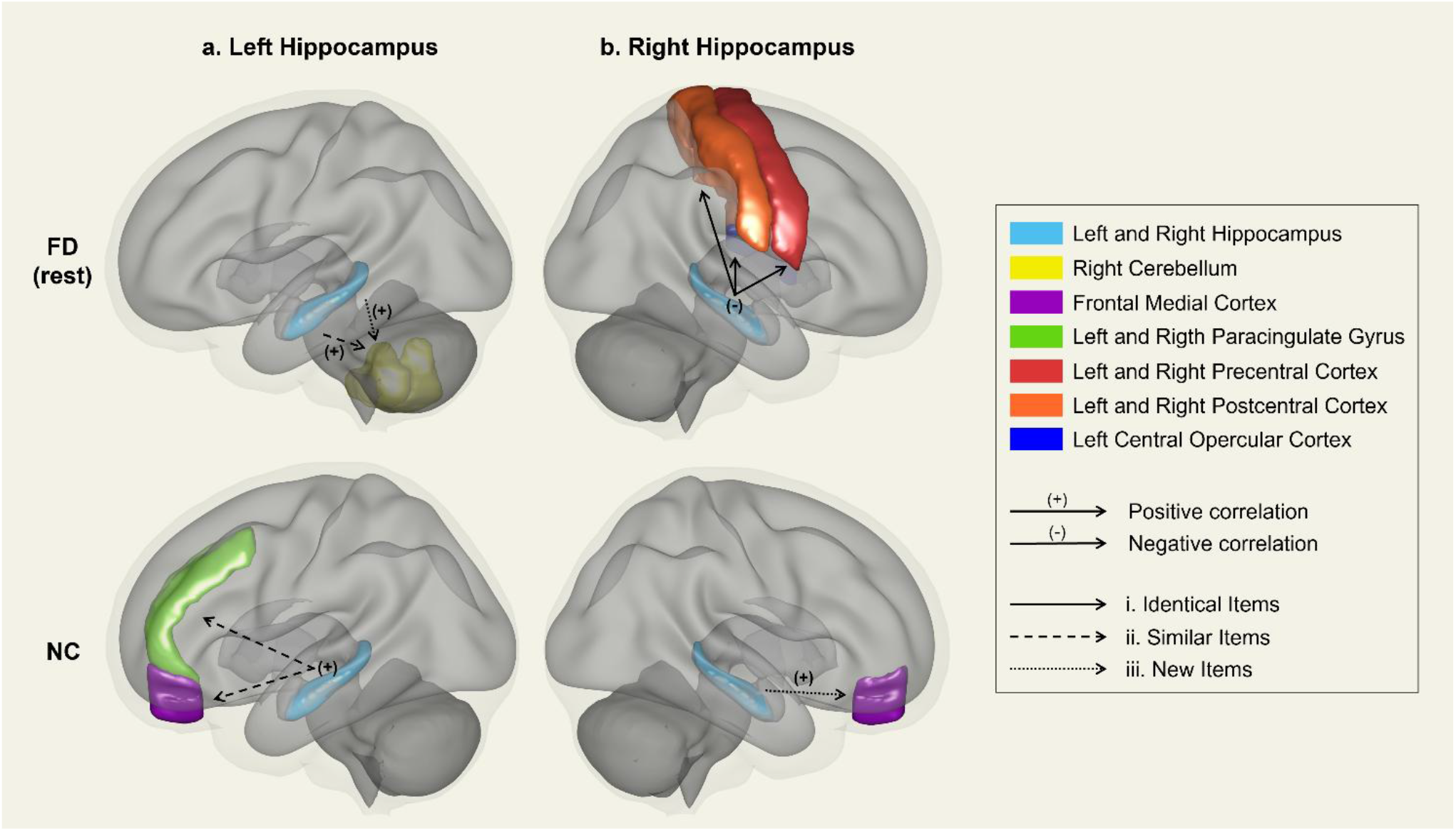
Correlations result between left and right hippocampal FC across conditions (NC and FD during rest after training) and episodic memory scores: GLM with FDR correction using CONN toolbox on MATLAB. a. Left hippocampus b. Right hippocampus. FD: Freedivers, NC: Controls.

#### 3.2 Right hippocampus

##### FD group, rest after training

We found a significant negative correlation between EM scores of i. *identical* items and the FC of the right hippocampus with the left postcentral gyrus (*r* = -0.91, p<0.001, n = 17), the left precentral cortex and left central opercular cortex (*r* = - 0.85, p<0.01, n = 17), and the right pre and postcentral gyrus (*r* = -0.83, p = 0.011, n = 17). No other significant correlations were found in other conditions.

##### NC group

We found a significant positive correlation between EM scores of iii. *new items* and the FC of the right hippocampus with the frontal medial cortex (r = 0.83, p = 0.017, n = 20) (Figure 8).

## Discussion

This study investigated functional connectivity changes associated with intermittent voluntary hypoxia in freedivers using a longitudinal design. Three main findings emerged: (1) large-scale network reorganization at rest after training, (2) selective hippocampal network reorganization before training that increased after training. (3) associations between hippocampal connectivity and episodic memory performance, particularly after training, suggesting compensatory neural adaptations to preserve cognitive abilities.

### 1. Whole-brain reorganization following freediving training

Whole-brain analyses revealed consistent modifications in connectivity across networks involved in cognitive control, attention, and sensorimotor processing, typically impaired under involuntary hypoxia (Micaux et al., 2025b). Increased connectivity was primarily observed between frontoparietal regions and sensorimotor, salience, and visual networks, with effects that became more pronounced after training.

In parallel, reduced connectivity was observed in occipital regions and in specific cerebellar-cortical interactions. Before training, hypoconnectivity during apnea mainly involved the occipital regions and the default mode network, ventral attention network, salience network, frontoparietal network, and cerebellum. After training, this pattern shifted toward reduced connectivity between the cerebellum and frontoparietal regions. This evolution suggests a reorganization process in which sensory-driven processing becomes less dominant, while internally oriented control networks become more prominent.

The involvement of the supplementary motor area (SMA) is consistent with the functional demands of freediving, which requires sustained motor inhibition and voluntary control of respiration (Allinger et al., 2024). Overall, these findings point to a reconfiguration of large-scale networks toward efficient coordination of cognitive control and sensorimotor regulation under repeated hypoxic exposure.

### 2. Hippocampal reorganization and functional adaptation following freediving training

The hippocampus showed a distinct and consistent pattern of functional reorganization. We observed consistent hypoconnectivity between the left hippocampus and the sensorimotor cortex across all conditions, more pronounced after training, along with decreased coupling to the default mode network. Simultaneously, hippocampal hyperconnectivity with the cerebellum and visual regions increased after training. This pattern suggests a shift in hippocampal network integration away from externally driven processes toward internally oriented and integrative systems. Reduced hippocampal-sensorimotor coupling may reflect decreased interference from sensory and motor signals, consistent with a shift toward more automatic processing, while increased hippocampal-cerebellar connectivity may support coordination between cognitive and physiological processes.

The cerebellum is increasingly recognized for its contribution to cognitive functions, including memory, through its interactions with hippocampal and cortical networks. Strengthened hippocampal-cerebellar connectivity may therefore facilitate integration across memory, prediction, and autonomic regulation systems, particularly under physiological constraints such as hypoxia.

Importantly, hippocampal changes differed from whole-brain trends. While large-scale networks showed increased integration involving sensorimotor and control systems, the hippocampus showed reduced coupling with these regions. This dissociation suggests that hippocampal reorganization follows a distinct functional trajectory, potentially prioritizing processes related to memory and internal representation.

These findings align with those of Annen et al. (2021), who reported strengthened regulatory and introspective networks in elite freedivers, similar to meditation, but specifically associated with the physiological demands of apnea. Our data extend these findings at the group level, revealing how hippocampal and cerebellar dynamics contribute to sustaining cognitive function during intermittent voluntary hypoxia.

Complementary to our results, prior research in exercise training (with hypoxia, as in synchronized swimming, or without), and low-dose hypoxia found similar effects, each offering insight into different mechanisms that may contribute to memory preservation in FD. Jones & Cooper, (2018) showed preserved oxygenation in the prefrontal cortex during maximal apneas in synchronized swimmers, despite severe muscle deoxygenation, indicating prioritized cerebral protection. Endurance training is known to promote brain efficiency through reduced global FC in non-essential regions and selective strengthening of networks relevant for a specific task (Li et al., 2023; Raichlen et al., 2016), similar to what we observed in our FD group. However, this is not unique to endurance exercise. For instance, research on skeleton athletes illustrates a clear case of specialization: increased rs-FC in regions involved in cognitive and motor control, reward learning, visual processing, spatial cognition, and emotional regulation, all of which are essential in their discipline. A distinct line of research on intermittent hypoxic training (IHT) suggests that low-dose hypoxia, when paired with exercise, could have neuroprotective effects (boost production of BDNF), and a beneficial impact on cognition (Boulares et al., 2024; Damgaard et al., 2023; Rybnikova et al., 2022). Our results support this: hippocampal FC patterns became more predictive of cognitive performance only after training, suggesting that hypoxia alone is not sufficient; adaptation depends on repeated, controlled exposure combined with exercise training. Not surprisingly, the impact of IHT depends on hypoxia intensity, with moderate levels promoting beneficial effects while more severe exposure leads to detrimental outcomes (Damgaard et al., 2023; Navarrete-Opazo and Mitchell, 2014).

Overall, neuroplasticity induced by freediving appears to reflect a unique convergence of sport and hypoxia adaptation. This combination led to a functional reorganization that prioritizes internal regulation, memory preservation, and network efficiency. Our data suggests, therefore, that under controlled and repeated exposure, voluntary hypoxia may support neural resilience. Notably, the cerebellum, often overlooked in rs-fMRI studies, emerged as a central adaptive hub connected to the hippocampus.

### 3. Hippocampal FC reflects episodic memory scores

After training, hippocampal functional connectivity was associated with episodic memory performance in freedivers, particularly for pattern separation-related measures. Positive associations between left hippocampal–cerebellar connectivity and memory performance suggest that this network interaction may support efficient discrimination of similar memory traces.

A shift was also observed in the neural substrates of recognition (i.e. new items): while control participants relied on right hippocampal–medial prefrontal pathways, FD increasingly engaged left hippocampal–cerebellar circuits. This suggests a redistributed encoding strategy following repeated hypoxia exposure. The negative correlation between right hippocampal-sensorimotor FC and identical items further indicates that sensorimotor disengagement may reduce interference and support accurate recognition; consistent with Simons & Spiers, (2003), who proposed that disengagement of irrelevant networks can facilitate memory by preventing information overload or cognitive noise. Notably, the pattern of associations differed between freedivers and controls. While controls showed hippocampal–prefrontal associations, freedivers exhibited stronger hippocampal– cerebellar relationships after training. This shift suggests that repeated hypoxic exposure may be associated with a redistribution of functional pathways supporting memory performance.

Taken together, these results highlight that hippocampal FC is related to episodic memory performance in FD, particularly after training. These findings align with Maeshima et al. (2017) who reported beneficial effects of regular synchronized swimming practice on memory, suggesting that freediving training may similarly lead to long-term cognitive improvements. Rather than reflecting a global enhancement of cortical connectivity (Amer and Davachi, 2023; Stevenson et al., 2020), this relationship appears to rely on a reorganization of network interactions involving both cortical and subcortical regions.

### 4. Future directions and perspectives

Our findings support the idea that voluntary intermittent hypoxia, when combined with physical training, is associated with functional brain reorganization. The hippocampus appears to be a key node in this process, with connectivity patterns that relate to cognitive performance. These observations are consistent with frameworks of adaptive neuroplasticity and cognitive reserve, in which functional reorganization contributes to maintaining performance under physiological constraints (Barulli and Stern, 2013; Verkhratsky and Zorec, 2024).

The longitudinal design is a notable strength of this study, as freedivers served as their own controls, enabling differentiation between baseline characteristics and changes observed after the training period.

Several limitations should nonetheless be acknowledged. The sample size was modest, and participants were exclusively male, a demographic composition that, while limiting generalizability, reflects the current profile of the freediving population. Future studies should aim to include larger and more diverse cohorts, particularly female athletes (Jung et al., 2020), as the sport continues to diversify. Additionally, multimodal imaging approaches would help further elucidate the physiological mechanisms underlying these functional changes. Designs including control groups with repeated imaging would be particularly valuable for clarifying causal relationships. Prolonged studies are also needed to assess the persistence and reversibility of these adaptations (Blackmore et al., 2024). Additionally, extending investigations to other cognitive domains may help determine whether these adaptations are specific to memory or reflect broader changes in brain function (McKeown et al., 2025). An additional methodological point concerns the fMRI paradigm: rather than a conventional resting-state acquisition, we combined the naturalistic Inscapes film (shown to preserve intrinsic network organization; Vanderwal et al., 2015) with apnea instructions. However, we do not expect this hybrid design to have substantially influenced the group differences reported here, since the film was identical across groups. This design strengthens our freediver-control comparison, as it helps ensure that the observed connectivity differences reflect apnea itself rather than differences in resting-state compliance. Future studies pairing conventional resting-state acquisitions with physiological measures (e.g., end-tidal CO2, cerebral blood flow) could further help disentangle neural adaptation from physiological state effects.

## Conclusion

Freediving training is associated with selective reorganization of hippocampal and large-scale brain networks, distinct from the widespread disruptions seen under involuntary hypoxia. These changes are linked to episodic memory performance and may reflect adaptive neuroplastic processes under repeated voluntary hypoxia. Freediving therefore provides a valuable human model for investigating functional brain adaptation and may inform therapeutic interventions to enhance cognitive resilience. Beyond sport, these insights open translational avenues for therapeutic interventions targeting hippocampal vulnerability, such as in aging, neurodegeneration, or hypoxia-related pathologies, through controlled hypoxic training paradigms designed to harness adaptive neuroplasticity.

## Acknowledgment

We express our sincere gratitude to the MRI technologists and the entire medical and nursing staff for their assistance throughout all stages of data acquisition. We are especially grateful to Mrs Mediouni and Mrs. Carré for their exceptional dedication and attentive care provided to the participants.

## REFERENCES

Allinger, J., Noulhiane, M., Féménias, D., Louvet, B., Clua, E., Bouyeure, A., Lemaître, F., 2024. Risk profiles of elite breath-hold divers. Int J Environ Health Res 1–13. 10.1080/09603123.2024.2368718

Amer, T., Davachi, L., 2023. Extra-hippocampal contributions to pattern separation. eLife 12, e82250. 10.7554/eLife.82250

Annen, J., Panda, R., Martial, C., Piarulli, A., Nery, G., Sanz, L.R.D., Valdivia-Valdivia, J.M., Ledoux, D., Gosseries, O., Laureys, S., 2021. Mapping the functional brain state of a world champion freediver in static dry apnea. Brain Struct Funct 226, 2675–2688. 10.1007/s00429-021-02361-1

Barulli, D., Stern, Y., 2013. Efficiency, capacity, compensation, maintenance, plasticity: emerging concepts in cognitive reserve. Trends Cogn Sci 17, 502–509. 10.1016/j.tics.2013.08.012

Blackmore, D.G., Schaumberg, M., Ziaei, M., Belford, S., To, X., O’Keeffe, I., Bernard, A., Mitchell, J., Hume, E., Rose, G.L., Shaw, T., York, A., Barth, M., Cooper, E.J., Skinner, T.L., Nasrallah, F.A., Riek, S., Bartlett, P.F., 2024. Long-Term Improvement in Hippocampal-Dependent Learning Ability in Healthy, Aged Individuals Following High Intensity Interval Training. Aging and Disease 16, 1732–1754. 10.14336/AD.2024.0642

Boulares, A., Pichon, A., Faucher, C., Bragazzi, N.L., Dupuy, O., 2024. Effects of Intermittent Hypoxia Protocols on Cognitive Performance and Brain Health in Older Adults Across Cognitive States: A Systematic Literature Review. J Alzheimers Dis 101, 13–30. 10.3233/JAD-240711

Castillon, C., Lunion, S., Desvignes, N., Hanauer, A., Laroche, S., Poirier, R., 2018. Selective alteration of adult hippocampal neurogenesis and impaired spatial pattern separation performance in the RSK2-deficient mouse model of Coffin-Lowry syndrome. Neurobiology of Disease 115, 69–81. 10.1016/j.nbd.2018.04.007

Damgaard, V., Mariegaard, J., Lindhardsen, J.M., Ehrenreich, H., Miskowiak, K.W., 2023. Neuroprotective Effects of Moderate Hypoxia: A Systematic Review. Brain Sciences 13, 1648. 10.3390/brainsci13121648

Heller-Wight, A., Phipps, C., Sexton, J., Ramirez, M., Warren, D.E., 2023. Hippocampal Resting State Functional Connectivity Associated with Physical Activity in Periadolescent Children. Brain Sci 13, 1558. 10.3390/brainsci13111558

Jones, B., Cooper, C.E., 2018. Near Infrared Spectroscopy (NIRS) Observation of Vastus Lateralis (Muscle) and Prefrontal Cortex (Brain) Tissue Oxygenation During Synchronised Swimming Routines in Elite Athletes. Adv Exp Med Biol 1072, 111–117. 10.1007/978-3-319-91287-5_18

Jung, M., Zou, L., Yu, J.J., Ryu, S., Kong, Z., Yang, L., Kang, M., Lin, J., Li, H., Smith, L., Loprinzi, P.D., 2020. Does exercise have a protective effect on cognitive function under hypoxia? A systematic review with meta-analysis. J Sport Health Sci 9, 562–577. 10.1016/j.jshs.2020.04.004

Khuu, M.A., Pagan, C.M., Nallamothu, T., Hevner, R.F., Hodge, R.D., Ramirez, J.-M., Garcia, A.J., 2019. Intermittent Hypoxia Disrupts Adult Neurogenesis and Synaptic Plasticity in the Dentate Gyrus. J Neurosci 39, 1320–1331. 10.1523/JNEUROSCI.1359-18.2018

Lemaître, F., 2015. L’apnée. De la théorie à la pratique. Presses universitaires de Rouen et du Havre.

Li, J., Fei, G.-H., 2013. The unique alterations of hippocampus and cognitive impairment in chronic obstructive pulmonary disease. Respir Res 14, 140. 10.1186/1465-9921-14-140

Li, W., Zhang, Q., Yang, R., Liu, B., Chen, G., Wang, B., Xu, T., Chen, J., Zhou, X., Wen, S., 2023. Characteristics of resting state functional connectivity of motor cortex of high fitness level college students: Experimental evidence from functional near infrared spectroscopy (fNIRS). Brain Behav 13, e3099. 10.1002/brb3.3099

Lu, G., Rili, G., Shuang, M., 2025. Impact of hypoxia on the hippocampus: A review. Medicine 104, e41479. 10.1097/MD.0000000000041479

Maeshima, E., Okumura, Y., Tatsumi, J., Tomokane, S., Ikeshima, A., 2017. Cognitive function in middle-aged and older adults participating in synchronized swimming-exercise. J Phys Ther Sci 29, 148–151. 10.1589/jpts.29.148

McKeown, D.J., Angus, D.J., Moustafa, A.A., Schinazi, V.R., 2025. Hypoxia and Cognitive Ability in Humans: A Systematic Review and Meta-Analysis. 10.1101/2025.05.15.654374

Micaux, J., Poiret, C., Zhao, J., El Hajj, A., Tillenon, M., Troudi Habibi, A., Mauconduit, F., Boumezbeur, F., Chiron, C., Noulhiane, M., 2025a. Does Freediving Lead to Hippocampal Adaptability to Hypoxia and Maintenance of Episodic Memory? J Integr Neurosci 24, 36672. 10.31083/JIN36672

Micaux, J., Troudi Habibi, A., Mauconduit, F., Noulhiane, M., 2025b. Hypoxia’s Impact on Hippocampal Functional Connectivity: Insights from Resting-State fMRI Studies. Brain Sci 15, 643. 10.3390/brainsci15060643

Navarrete-Opazo, A., Mitchell, G.S., 2014. Therapeutic potential of intermittent hypoxia: a matter of dose. Am J Physiol Regul Integr Comp Physiol 307, R1181–1197. 10.1152/ajpregu.00208.2014

Olaithe, M., Bucks, R.S., Hillman, D.R., Eastwood, P.R., 2018. Cognitive deficits in obstructive sleep apnea: Insights from a meta-review and comparison with deficits observed in COPD, insomnia, and sleep deprivation. Sleep Med Rev 38, 39–49. 10.1016/j.smrv.2017.03.005

Pfister, K.M., Stoyell, S.M., Miller, Z.R., Hunt, R.H., Zorn, E.P., Thomas, K.M., 2023. Reduced Hippocampal Volumes in Children with History of Hypoxic Ischemic Encephalopathy after Therapeutic Hypothermia. Children (Basel) 10, 1005. 10.3390/children10061005

Raichlen, D.A., Bharadwaj, P.K., Fitzhugh, M.C., Haws, K.A., Torre, G.-A., Trouard, T.P., Alexander, G.E., 2016. Differences in Resting State Functional Connectivity between Young Adult Endurance Athletes and Healthy Controls. Front Hum Neurosci 10, 610. 10.3389/fnhum.2016.0610

Ridgway, L., McFarland, K., 2006. Apnea diving: long-term neurocognitive sequelae of repeated hypoxemia. Clin Neuropsychol 20, 160–176. 10.1080/13854040590947407

Rybnikova, E.A., Nalivaeva, N.N., Zenko, M.Y., Baranova, K.A., 2022. Intermittent Hypoxic Training as an Effective Tool for Increasing the Adaptive Potential, Endurance and Working Capacity of the Brain. Front Neurosci 16, 941740. 10.3389/fnins.2022.941740

Scoville, W.B., Milner, B., 1957. Loss of recent memory after bilateral hippocampal lesions. J Neurol Neurosurg Psychiatry 20, 11–21. 10.1136/jnnp.20.1.11

Simons, J.S., Spiers, H.J., 2003. Prefrontal and medial temporal lobe interactions in long-term memory. Nat Rev Neurosci 4, 637–648. 10.1038/nrn1178

Song, X., Roy, B., Kang, D.W., Aysola, R.S., Macey, P.M., Woo, M.A., Yan-Go, F.L., Harper, R.M., Kumar, R., 2018. Altered resting-state hippocampal and caudate functional networks in patients with obstructive sleep apnea. Brain Behav 8, e00994. 10.1002/brb3.994

Spallazzi, M., Dobisch, L., Becke, A., Berron, D., Stucht, D., Oeltze-Jafra, S., Caffarra, P., Speck, O., Düzel, E., 2019. Hippocampal vascularization patterns: A high-resolution 7 Tesla time-of-flight magnetic resonance angiography study. Neuroimage Clin 21, 101609. 10.1016/j.nicl.2018.11.019

Stevenson, R.F., Reagh, Z.M., Chun, A.P., Murray, E.A., Yassa, M.A., 2020. Pattern Separation and Source Memory Engage Distinct Hippocampal and Neocortical Regions during Retrieval. J Neurosci 40, 843–851. 10.1523/JNEUROSCI.0564-19.2019

Suwabe, K., Byun, K., Hyodo, K., Reagh, Z.M., Roberts, J.M., Matsushita, A., Saotome, K., Ochi, G., Fukuie, T., Suzuki, K., Sankai, Y., Yassa, M.A., Soya, H., 2018. Rapid stimulation of human dentate gyrus function with acute mild exercise. Proc Natl Acad Sci U S A 115, 10487–10492. 10.1073/pnas.1805668115

Vanderwal, T., Kelly, C., Eilbott, J., Mayes, L.C., Castellanos, F.X., 2015. Inscapes: A movie paradigm to improve compliance in functional magnetic resonance imaging. NeuroImage 122, 222–232. 10.1016/j.neuroimage.2015.07.069

Verkhratsky, A., Zorec, R., 2024. Neuroglia in cognitive reserve. Mol Psychiatry 29, 3962–3967. 10.1038/s41380-024-02644-z

Wang, X., Cui, L., Ji, X., 2022. Cognitive impairment caused by hypoxia: from clinical evidences to molecular mechanisms. Metab Brain Dis 37, 51–66. 10.1007/s11011-021-00796-3

Zhang, Z.-A., Sun, Y., Yuan, Z., Wang, L., Dong, Q., Zhou, Y., Zheng, G., Aschner, M., Zou, Y., Luo, W., 2022. Insight into the Effects of High-Altitude Hypoxic Exposure on Learning and Memory. Oxid Med Cell Longev 2022, 4163188. 10.1155/2022/4163188

Zhao, J., Noulhiane, M., In Prep. An ecological tool evaluating episodic memory patterns (Pattern Separation and Pattern Completion).

