## Supplementary_material for "Hippocampal and large-scale functional connectivity reorganization following freediving training, and relationships with episodic memory"

**Table S1**: Significant p-values of seed-to-voxel comparison (hyperconnectivity) with FDR correction. AD: Anterior Division; PD: Posterior Division. FD: Freedivers, NC, Controls.

| Seed Region | Comparison | Hyperconnectivity | p-FDR |
|---|---|---|---|
| Hippocampus_Left | $FDt_{1apnea}$ > NC | Left & Right Cerebellum | < 0.001 |
| Hippocampus_Left | $FDt_{1apnea}$ > NC | Right Temporal Occipital Fusiform Cortex | < 0.001 |
| Hippocampus_Left | $FDt_{1apnea}$ > NC | Right Temporal Fusiform Cortex | < 0.001 |
| Hippocampus_Left | $FDt_{0rest}$ > NC | Left Cerebellum | 0.009 |
| Hippocampus_Left | $FDt_{1rest}$ > NC | Left & Right Cerebellum | 0.003; 0.01 |
| Hippocampus_Right | $FDt_{0apnea}$ > NC | Right Cerebellum | < 0.001 |
| Hippocampus_Right | $FDt_{1apnea}$ > NC | Left & Right Cerebellum | < 0.001 |
| Hippocampus_Right | $FDt_{1rest}$ > NC | Right Cerebellum | 0.026 |
| FP_Left | $FDt_{1apnea}$ > $FDt_{1rest}$ | Left & Right Intracalcarine Cortex | < 0.001; 0.003 |
| FP_Left | $FDt_{1apnea}$ > $FDt_{1rest}$ | Right Cuneal Cortex | < 0.001 |
| FP_Left | $FDt_{1apnea}$ > $FDt_{1rest}$ | Left & Right Lingual Gyrus | < 0.001; 0.003 |
| FP_Left | $FDt_{0apnea}$ > NC | Left & Right Central Opercular Cortex | 0.016 |
| FP_Left | $FDt_{0apnea}$ > NC | Left Postcentral Gyrus | 0.016 |
| FP_Left | $FDt_{0apnea}$ > NC | Right Precentral Gyrus | 0.028 |
| FP_Left | $FDt_{1apnea}$ > NC | Left Precentral Gyrus | 0.048 |
| FP_Left | $FDt_{1apnea}$ > NC | Right Postcentral Gyrus | 0.048 |
| FP_Left | $FDt_{1rest}$ > NC | Right Frontal Pole | 0.044 |
| FP_Left | $FDt_{1rest}$ > NC | Right Middle Frontal Gyrus | 0.044 |
| FP_Left | $FDt_{1rest}$ > NC | Right Superior Frontal Gyrus | 0.044 |
| FP_Right | $FDt_{0apnea}$ > NC | Left & Right Precentral Gyrus | < 0.001 |
| FP_Right | $FDt_{0apnea}$ > NC | Left & Right Postcentral Gyrus | < 0.001 |
| FP_Right | $FDt_{0apnea}$ > NC | Left & Right Central Opercular Cortex | < 0.001 |
| FP_Right | $FDt_{0apnea}$ > NC | Left Superior Temporal Gyrus | < 0.001 |
| FP_Right | $FDt_{0apnea}$ > NC | Left Middle Temporal Gyrus | < 0.001 |
| FP_Right | $FDt_{0apnea}$ > NC | Left Planum Temporal | < 0.001 |
| FP_Right | $FDt_{0apnea}$ > NC | Right Insular Cortex | < 0.001 |
| FP_Right | $FDt_{0apnea}$ > NC | Right Planum Polare | < 0.001 |
| FP_Right | $FDt_{1apnea}$ > NC | Left & Right Precentral Gyrus | < 0.001 |
| FP_Right | $FDt_{1apnea}$ > NC | Left & Right Postcentral Gyrus | < 0.001 |
| FP_Right | $FDt_{1apnea}$ > NC | Left Middle Frontal Gyrus | < 0.001 |
| FP_Right | $FDt_{1apnea}$ > NC | Left Inferior Temporal Gyrus | 0.041 |
| FP_Right | $FDt_{1apnea}$ > NC | Right Insular Cortex | 0.047 |
| FP_Right | $FDt_{1apnea}$ > NC | Right Central Opercular Cortex | 0.047 |

| | | | |
|---|---|---|---|
| FP_Right | $FDt_{1apnea}$ > NC | Right Parietal Operculum Cortex | 0.047 |
| FP_Right | $FDt_{0rest}$ > NC | Left Inferior Frontal Gyrus | 0.01 |
| FP_Right | $FDt_{0rest}$ > NC | Left Frontal Orbital Cortex | 0.01 |
| FP_Right | $FDt_{1rest}$ > NC | Left & Right Precentral Gyrus | < 0.001 |
| FP_Right | $FDt_{1rest}$ > NC | Left & Right Postcentral Gyrus | < 0.001 |
| FP_Right | $FDt_{1rest}$ > NC | Left Inferior Temporal Gyrus | 0.049 |
| FPN_Left | $FDt_{0apnea}$ > NC | Right Precentral Gyrus | 0.006 |
| FPN_Left | $FDt_{1apnea}$ > NC | Left & Right Frontal Pole | < 0.001 |
| FPN_Left | $FDt_{1apnea}$ > NC | Left & Right Paracingulate Gyrus | < 0.001 |
| FPN_Left | $FDt_{1apnea}$ > NC | Cingulate Gyrus AD | < 0.001 |
| FPN_Left | $FDt_{1apnea}$ > NC | Left Postcentral Gyrus | 0.039 |
| FPN_Left | $FDt_{1apnea}$ > NC | Cingulate Gyrus PD | 0.039 |
| FPN_Left | $FDt_{0rest}$ > NC | Cingulate Gyrus PD | 0.003 |
| FPN_Left | $FDt_{0rest}$ > NC | Left & Right Paracingulate Gyrus | 0.032; 0.049 |
| FPN_Right | $FDt_{0apnea}$ > NC | Left & Right Precentral Gyrus | < 0.001 |
| FPN_Right | $FDt_{0apnea}$ > NC | Left & Right Postcentral Gyrus | < 0.001 |
| FPN_Right | $FDt_{0apnea}$ > NC | Right Superior Parietal Lobule | < 0.001 |
| FPN_Right | $FDt_{0apnea}$ > NC | Left & Right Supplementary Motor Area | < 0.001 |
| FPN_Right | $FDt_{0apnea}$ > NC | Cingulate Gyrus AD | < 0.001 |
| FPN_Right | $FDt_{0apnea}$ > NC | Left Planum Temporal | 0.006 |
| FPN_Right | $FDt_{0apnea}$ > NC | Left Heschl's Gyrus | 0.006 |
| FPN_Right | $FDt_{1apnea}$ > NC | Left & Right Precentral Gyrus | < 0.001 |
| FPN_Right | $FDt_{1apnea}$ > NC | Left & Right Postcentral Gyrus | < 0.001 |
| FPN_Right | $FDt_{1apnea}$ > NC | Left Frontal Pole | 0.003 |
| FPN_Right | $FDt_{1apnea}$ > NC | Left & Right Central Opercular Cortex | 0.005; 0.008 |
| FPN_Right | $FDt_{1apnea}$ > NC | Left & Right Parietal Operculum Cortex | 0.005; 0.008 |
| FPN_Right | $FDt_{1apnea}$ > NC | Left & Right Insular Cortex | 0.005; 0.008 |
| FPN_Right | $FDt_{1apnea}$ > NC | Left & Right Heschl's Gyrus | 0.005; 0.008 |
| FPN_Right | $FDt_{1apnea}$ > NC | Left Planum Temporal | 0.008 |
| FPN_Right | $FDt_{1apnea}$ > NC | Left Paracingulate Gyrus | 0.022 |
| FPN_Right | $FDt_{0rest}$ > NC | Left & Right Precentral Gyrus | < 0.001 |
| FPN_Right | $FDt_{0rest}$ > NC | Left & Right Postcentral Gyrus | < 0.001 |
| FPN_Right | $FDt_{0rest}$ > NC | Left Occipital Pole | < 0.001 |
| FPN_Right | $FDt_{0rest}$ > NC | Left Lateral Occipital Cortex | < 0.001 |
| FPN_Right | $FDt_{0rest}$ > NC | Left Inferior Frontal Gyrus | < 0.001 |
| FPN_Right | $FDt_{0rest}$ > NC | Left Frontal Operculum Cortex | < 0.001 |
| FPN_Right | $FDt_{0rest}$ > NC | Left Paracingulate Gyrus | 0.007 |

| | | | |
|---|---|---|---|
| FPN_Right | $FDt_{0rest}$ > NC | Left Middle Temporal Gyrus | 0.028 |
| FPN_Right | $FDt_{0rest}$ > NC | Left Superior Temporal Gyrus | 0.028 |
| FPN_Right | $FDt_{1rest}$ > NC | Left & Right Precentral Gyrus | < 0.001; 0.037 |
| FPN_Right | $FDt_{1rest}$ > NC | Right Postcentral Gyrus | < 0.001 |
| OP_Left | $FDt_{0rest}$ > NC | Left & Right Middle Temporal Gyrus | < 0.001 |
| OP_Left | $FDt_{0rest}$ > NC | Left & Right Inferior Temporal Gyrus | < 0.001 |
| OP_Left | $FDt_{0rest}$ > NC | Left & Right Superior Temporal Gyrus | < 0.001 |
| OP_Left | $FDt_{0rest}$ > NC | Left & Right Angular Gyrus | < 0.001 |
| OP_Left | $FDt_{0rest}$ > NC | Left & Right Supramarginal Gyrus | < 0.001 |
| OP_Left | $FDt_{0rest}$ > NC | Left & Right Superior Parietal Lobule | < 0.001 |
| OP_Left | $FDt_{0rest}$ > NC | Left & Right Lateral Occipital Cortex | < 0.001; 0.03 |
| OP_Left | $FDt_{0rest}$ > NC | Right Middle Frontal Gyrus | < 0.001 |
| OP_Right | $FDt_{0apnea}$ > NC | Left & Right Precentral Gyrus | < 0.001 |
| OP_Right | $FDt_{0apnea}$ > NC | Left & Right Superior Frontal Gyrus | < 0.001 |
| OP_Right | $FDt_{0apnea}$ > NC | Left Middle Frontal Gyrus | < 0.001 |
| OP_Right | $FDt_{0apnea}$ > NC | Left Supplementary Motor Area | < 0.001 |
| OP_Right | $FDt_{0rest}$ > NC | Left & Right Middle Temporal Gyrus | < 0.001 |
| OP_Right | $FDt_{0rest}$ > NC | Left & Right Inferior Temporal Gyrus | < 0.001 |
| OP_Right | $FDt_{0rest}$ > NC | Left & Right Superior Temporal Gyrus | < 0.001 |
| OP_Right | $FDt_{0rest}$ > NC | Left & Right Angular Gyrus | < 0.001; 0.001 |
| OP_Right | $FDt_{0rest}$ > NC | Left & Right Supramarginal Gyrus | < 0.001; 0.001 |
| OP_Right | $FDt_{0rest}$ > NC | Left & Right Superior Parietal Lobule | < 0.001; 0.001 |
| OP_Right | $FDt_{0rest}$ > NC | Right Middle Frontal Gyrus | 0.014 |

**Table S2**: Significant p-values of seed-to-voxel comparison (hypoconnectivity) with FDR correction. AD: Anterior Division; PD: Posterior Division. FD: Freedivers, NC, Controls.

| Seed Region | Comparison | Hypoconnectivity | p-FDR |
|---|---|---|---|
| DMN | $FDt_{1apnea}$ > $FDt_{1rest}$ | Left & Right Lateral Occipital Cortex | 0.034 |
| DMN | $FDt_{1apnea}$ > $FDt_{1rest}$ | Left Occipital Pole | 0.034 |
| DMN | $FDt_{1apnea}$ > NC | Left Occipital Pole | < 0.001 |
| DMN | $FDt_{1apnea}$ > NC | Left Intracalcarine Cortex | < 0.001 |
| DMN | $FDt_{1apnea}$ > NC | Left Supracalcarine Cortex | < 0.001 |
| DMN | $FDt_{1apnea}$ > NC | Right Lingual Gyrus | 0.007 |
| DMN | $FDt_{1apnea}$ > NC | Right Occipital Fusiform Gyrus | 0.007 |
| DMN | $FDt_{1apnea}$ > NC | Right Cerebellum | 0.007 |
| Hippocampus_Left | $FDt_{0apnea}$ > NC | Left & Right Precentral Gyrus | < 0.001 |
| Hippocampus_Left | $FDt_{0apnea}$ > NC | Left & Right Postcentral Gyrus | < 0.001 |
| Hippocampus_Left | $FDt_{0apnea}$ > NC | Left & Right Supplementary Motor Area | < 0.001 |

| | | | |
|---|---|---|---|
| Hippocampus_Left | $FDt_{0apnea}$ > NC | Right Superior Parietal Lobule | < 0.001 |
| Hippocampus_Left | $FDt_{0apnea}$ > NC | Left Paracingulate Gyrus | < 0.001 |
| Hippocampus_Left | $FDt_{1apnea}$ > NC | Left & Right Precentral Gyrus | < 0.001 |
| Hippocampus_Left | $FDt_{1apnea}$ > NC | Left & Right Postcentral Gyrus | < 0.001 |
| Hippocampus_Left | $FDt_{1apnea}$ > NC | Left & Right Supplementary Motor Area | < 0.001 |
| Hippocampus_Left | $FDt_{0rest}$ > NC | Left & Right Temporal Pole | 0.009; 0.047 |
| Hippocampus_Left | $FDt_{0rest}$ > NC | Right Middle Temporal Gyrus | 0.009 |
| Hippocampus_Left | $FDt_{0rest}$ > NC | Frontal Medial Cortex | 0.047 |
| Hippocampus_Left | $FDt_{1rest}$ > NC | Frontal Medial Cortex | < 0.001 |
| Hippocampus_Left | $FDt_{1rest}$ > NC | Left Precentral Gyrus | 0.041 |
| Hippocampus_Left | $FDt_{1rest}$ > NC | Left & Right Postcentral Gyrus | 0.008; 0.006 |
| Hippocampus_Left | $FDt_{1rest}$ > NC | Left & Right Superior Parietal Lobule | 0.006 |
| Hippocampus_Left | $FDt_{1rest}$ > NC | Left & Right Superior Temporal Gyrus | 0.013; 0.018 |
| Hippocampus_Left | $FDt_{1rest}$ > NC | Left & Right Planum Polare | 0.037; 0.018 |
| Hippocampus_Left | $FDt_{1rest}$ > NC | Right Central Opercular Cortex | 0.018 |
| Hippocampus_Left | $FDt_{1rest}$ > NC | Precuneus Cortex | 0.041 |
| Hippocampus_Right | $FDt_{1apnea}$ > NC | Left Precentral Gyrus | 0.039 |
| Hippocampus_Right | $FDt_{1rest}$ > NC | Left Postcentral Gyrus | < 0.001 |
| Hippocampus_Right | $FDt_{1rest}$ > NC | Left Precentral Gyrus | < 0.001 |
| Hippocampus_Right | $FDt_{1rest}$ > NC | Right Planum Temporal | 0.046 |
| Hippocampus_Right | $FDt_{1rest}$ > NC | Right Planum Polare | 0.046 |
| Hippocampus_Right | $FDt_{1rest}$ > NC | Right Central Opercular Cortex | 0.046 |
| FP_Left | $FDt_{0rest}$ > NC | Right Cerebellum | < 0.001 |
| FP_Left | $FDt_{1rest}$ > NC | Right Intracalcarine Cortex | 0.001 |
| FP_Right | $FDt_{0rest}$ > NC | Left & Right Cerebellum | < 0.001 |
| FP_Right | $FDt_{1rest}$ > NC | Right Intracalcarine Cortex | 0.049 |
| FPN_Left | $FDt_{0rest}$ > NC | Left & Right Cerebellum | < 0.001; 0.045 |
| FPN_Left | $FDt_{0rest}$ > NC | Left Thalamus | 0.045 |
| FPN_Left | $FDt_{1rest}$ > NC | Right Intracalcarine Cortex | 0.044 |
| FPN_Right | $FDt_{1apnea}$ > NC | Left Caudate | 0.045 |
| FPN_Right | $FDt_{0rest}$ > NC | Left & Right Cerebellum | 0.024 |
| FPN_Right | $FDt_{1rest}$ > NC | Left Cerebellum | 0.004 |
| FPN_Right | $FDt_{1rest}$ > NC | Right Thalamus | 0.037 |
| OP_Left | $FDt_{0apnea}$ > $FDt_{0rest}$ | Left & Right Middle Temporal Gyrus | < 0.001; 0.001 |
| OP_Left | $FDt_{0apnea}$ > $FDt_{0rest}$ | Left Inferior Temporal Gyrus | < 0.001 |
| OP_Left | $FDt_{0apnea}$ > $FDt_{0rest}$ | Left Angular Gyrus | < 0.001 |
| OP_Left | $FDt_{0apnea}$ > $FDt_{0rest}$ | Left Lateral Occipital Cortex | < 0.001 |

| | | | |
|---|---|---|---|
| OP_Left | $FDt_{0apnea}$ > $FDt_{0rest}$ | Left Supramarginal Gyrus | < 0.001 |
| OP_Left | $FDt_{0apnea}$ > $FDt_{0rest}$ | Cingulate Gyrus PD | 0.001 |
| OP_Left | $FDt_{0apnea}$ > $FDt_{0rest}$ | Left Frontal Pole | 0.038 |
| OP_Left | $FDt_{0apnea}$ > $FDt_{0rest}$ | Left Paracingulate Gyrus | 0.038 |
| OP_Left | $FDt_{1apnea}$ > $FDt_{1rest}$ | Left Lateral Occipital Cortex | 0.005 |
| OP_Left | $FDt_{1apnea}$ > $FDt_{1rest}$ | Precuneus Cortex | 0.007 |
| OP_Left | $FDt_{1apnea}$ > $FDt_{1rest}$ | Cingulate Gyrus PD | 0.007 |
| OP_Left | $FDt_{0apnea}$ > NC | Brain Stem | 0.049 |
| OP_Left | $FDt_{0apnea}$ > NC | Frontal Medial Cortex | 0.049 |
| OP_Left | $FDt_{0apnea}$ > NC | Right Occipital Fusiform Gyrus | 0.049 |
| OP_Left | $FDt_{1apnea}$ > NC | Left Paracingulate Gyrus | 0.001 |
| OP_Left | $FDt_{1apnea}$ > NC | Cingulate Gyrus AD | 0.001 |
| OP_Left | $FDt_{0rest}$ > NC | Left & Right Lingual Gyrus | < 0.001 |
| OP_Left | $FDt_{0rest}$ > NC | Left & Right Occipital Fusiform Gyrus | < 0.001 |
| OP_Left | $FDt_{0rest}$ > NC | Left & Right Cerebellum | < 0.001 |
| OP_Left | $FDt_{0rest}$ > NC | Left & Right Occipital Pole | < 0.001 |
| OP_Left | $FDt_{0rest}$ > NC | Vermis | < 0.001 |
| OP_Left | $FDt_{0rest}$ > NC | Right Thalamus | < 0.001 |
| OP_Left | $FDt_{0rest}$ > NC | Brain Stem | < 0.001 |
| OP_Right | $FDt_{0apnea}$ > $FDt_{0rest}$ | Right Middle Temporal Gyrus | 0.047 |
| OP_Right | $FDt_{0apnea}$ > $FDt_{0rest}$ | Right Cerebellum | 0.047 |
| OP_Right | $FDt_{0apnea}$ > NC | Right Occipital Fusiform Gyrus | < 0.001 |
| OP_Right | $FDt_{0apnea}$ > NC | Right Lingual Gyrus | < 0.001 |
| OP_Right | $FDt_{0apnea}$ > NC | Right Occipital Pole | < 0.001 |
| OP_Right | $FDt_{0apnea}$ > NC | Right Cerebellum | < 0.001 |
| OP_Right | $FDt_{0apnea}$ > NC | Vermis | < 0.001 |
| OP_Right | $FDt_{0apnea}$ > NC | Brain Stem | 0.011 |
| OP_Right | $FDt_{0apnea}$ > NC | Left Putamen | 0.02 |
| OP_Right | $FDt_{0apnea}$ > NC | Left Insular Cortex | 0.02 |
| OP_Right | $FDt_{0rest}$ > NC | Left & Right Lingual Gyrus | < 0.001 |
| OP_Right | $FDt_{0rest}$ > NC | Left & Right Occipital Fusiform Gyrus | < 0.001 |
| OP_Right | $FDt_{0rest}$ > NC | Left Cerebellum | < 0.001 |
| OP_Right | $FDt_{0rest}$ > NC | Right Occipital Pole | < 0.001 |
| OP_Right | $FDt_{0rest}$ > NC | Vermis | < 0.001 |
| OP_Right | $FDt_{0rest}$ > NC | Right Thalamus | < 0.001 |
| OP_Right | $FDt_{0rest}$ > NC | Brain Stem | < 0.001 |

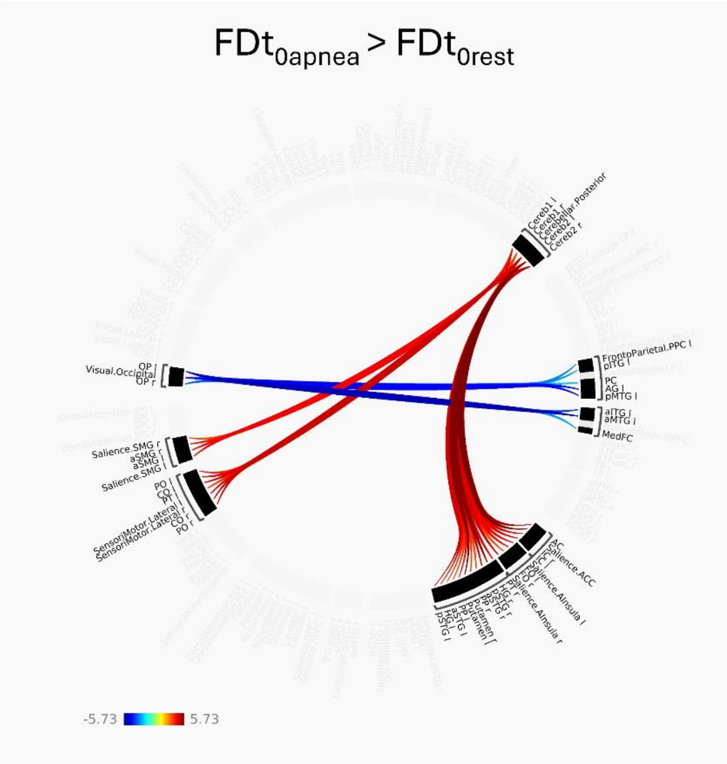


***Figure S1**: FC matrice showing differences in whole brain hypoconnectivity and hyperconnectivity comparing FDt$_{0apnea}$ > FDt$_{0rest}$. ROI-to-ROI analysis using CONN toolbox on MATLAB. N= 17 FD. FDR correction applied. FD: Freedivers.*
